# Effect of acetic acid bacteria colonization on oviposition and feeding site choice in *Drosophila suzukii* and its related species

**DOI:** 10.1101/2023.03.20.533419

**Authors:** Airi Sato, Joanne Y. Yew, Aya Takahashi

## Abstract

Oviposition site choice has a large impact on offspring performance. Unlike other vinegar flies that colonize decaying fruits, *Drosophila suzukii* lay eggs into hard ripening fruits by using their enlarged and serrated ovipositors (oviscapts). This behavior has an advantage over other species by providing access to the host fruit earlier and avoiding competition. However, the larvae are not fully adapted to a low-protein diet, and the availability of intact healthy fruits is seasonally restricted. Thus, to investigate oviposition site preference for microbial growth in this species, we conducted an oviposition assay using single species of commensal *Drosophila* acetic acid bacteria, *Acetobacter* and *Gluconobacter*. The oviposition site preferences for media with or without bacterial growth were quantified in multiple strains of *D. suzukii* and its closely related species, *D. subpulchrella* and *D. biarmipes*, and a typical fermenting-fruit consumer, *D. melanogaster*. Our comparisons demonstrated a continuous degree of preference for sites with *Acetobacter* growth both within and across species, suggesting that the niche separation is notable but not complete. The preference for *Gluconobacter* showed large variations among replicates and no clear differences between the strains. In addition, the lack of interspecific differences in feeding site preference for *Acetobacter*-containing media implies that the interspecific divergence in oviposition site preference occurred independently from the feeding site preference. Our oviposition assays measuring the preference of multiple strains from each fly species for acetic acid bacteria growth revealed intrinsic properties of shared resource usage among these fruit fly species.

## 1 Introduction

Fermenting fruits are nutrient-rich food resource for many insects including the larvae of various fruit flies. The flies consume a microbe-rich diet with an abundant supply of proteins necessary for the larval growth. Thus, the majority of *Drosophila* females lay eggs onto fermenting or rotting fruits. Whereas the females of *Drosophila suzukii*, the spotted wing drosophila, lay eggs into hard ripening fruits with relatively low P:C by using their enlarged and serrated ovipositors (oviscapts) (Walsh et al., 2011; Cini et al., 2012; Atallah et al., 2014). This behavior, which causes significant agricultural damage in recently invaded areas (Cini et al., 2012; Asplen et al., 2015), has allowed the offspring to have an advantage over other species by providing access to the host fruit earlier, thus avoiding competition.

However, considering that *D. suzukii* larvae have limited physiological adaptation to a low-protein diet and intact healthy fruits have seasonally restricted availability, the competitive advantage of ovipositing in ripening fruits can be conditional (Silva-Soares et al., 2017; Young et al., 2018; Kienzle et al., 2020; Deans and Hutchingson 2021). Therefore, the oviposition site preference towards non-fermenting fruits may depend on multiple factors, therefore, there is likely to be variability maintained within species. Also, since adult flies, especially females, require a large amount of protein for reproduction (Jensen et al., 2015), their foraging decisions are affected by their own nutritional demands as well (Lihoreau et al., 2016). Given the potential conflict between nutritional demand and competition for resources, we investigated the following: 1) the degree of interspecific differences and intraspecific variation in preference for oviposition sites that contain microbial species associated with decaying fruits, and 2) whether oviposition site preferences are independent from feeding site selection.

To pursue these questions, we conducted an oviposition assay using single species of *Drosophila* commensal acetic acid bacteria, *Acetobacter* and *Gluconobacter* (Broderick and Lemaitre 2012; Chandler et al., 2014; Vacchini et al., 2017). The oviposition site preferences for media with and without microbial growth were quantified in six strains of *D. suzukii*, in comparison to a typical fermenting fruit consumer, *D. melanogaster*. We also quantified the preferences of two strains from sibling species, *D. subpulchrella*, that has recently diverged from *D. suzukii*, and two strains from *D. biarmipes*, which is the most closely related species examined that prefer oviposition substrate colonized by microbes (Keesey et al., 2015; Sato et al., 2021). Our comparisons of the oviposition site preferences demonstrated a continuous degree of preference for microbes both within and across species, while the feeding assay indicated that microbial growth is not a factor that elicits interspecific differences in feeding site preferences among the tested species.

## 2 Materials and methods

### 2.1 Fly strains

The following strains were used to test the oviposition and feeding site preference for acetic acid bacteria: *D. suzukii* strain TMUS05 and TMUS08 collected in Hachioji, Japan, in 2015, *D. suzukii* strain Hilo collected in Hilo, Island of Hawai‘i, U. S. A., in 2017, *D. suzukii* strain OR collected in Oregon, U. S. A., in 2017, *D. suzukii* strain WT3 collected in California, U. S. A., in 2009 and sib-mated for ten generations (Chiu et al., 2013), *D. suzukii* strain YAM1 collected in Yamagata prefecture, Japan, in 2004, *D. subpulchrella* strain H243 collected in Hiratsuka, Japan, in 1979, *D. subpulchrella* strain M4 collected in Matsumoto, Japan, in 1982, *D. biarmipes* strain MYS118 collected in Mysore, India, in 1981, *D. biarmipes* strain NN68 collected in Nakhonn Nayok, Thailand, in 1977, and *D. melanogaster* strain Canton S BL#9515. *D. suzukii* and *D. subpulchrella* were maintained at 20 ± 1°C and other strains were maintained at 25 ± 1°C. All the strains were reared under the 12 h light: 12 h dark light condition. Flies were fed with standard corn meal food mixed with yeast, glucose, and agar. *D. suzukii* and *D. subpulchrella* flies aged 10–15 days after eclosion and *D. biarmipes* and *D. melanogaster* flies aged 4–7 days after eclosion were used for the assays.

### 2.2 Acetic acid bacteria

Single colonies of acetic acid bacteria were isolated from the microbes collected from the surface of fly-inoculated media and subjected to 16S-rRNA gene sequencing (Sato *et al*., 2021). The colonies of *Acetobacter* sp. and *Gluconobacter* sp. were identified by the 16S-rRNA gene sequences (Supplementary Data S1).

### 2.3 Oviposition assay to assess the preference for substrates with acetic acid bacteria

The oviposition assay was conducted in an oviposition chamber (90 mm diameter × 20 mm height petri dish, SH90-20, IWAKI) with test and control substrates. The substrates were made from 50% apple juice (SUNPACK, JAN code: 4571247510950) including 1% agar (Drosophila agar type II, Apex), and put in a petri dish (40 mm diameter × 13 mm height). Twenty μL of the bacterial solution (OD = 1 in distilled water) or the control distilled water were spread onto the surface of the substrate and incubated for 24 h at 25 ± 1°C.

Ten (for *D. suzukii* and *D. subpulchrella*) or 5 (for *D. melanogaster* and *D. biarmipes*) females were placed into each chamber without anesthesia by an aspirator within 4 h before the dark cycle and kept for 16 h under 12 h light: 12 h dark light conditions. The assay was conducted at 20 ± 1°C for *D. suzukii* and *D. subpulchrella* and at 25 ± 1°C for *D. biarmipes* and *D. melanogaster*. After the oviposition assay, photo images of each substrate with eggs were taken by a camera (Olympus OM-D E-M10 MarkII) with transmitted light from the bottom. The number of eggs on each substrate was counted using ImageJ v1.53k (Schneider et al., 2012).

The oviposition preference index (PI) for substrates inoculated with microbes was calculated using the following formula:

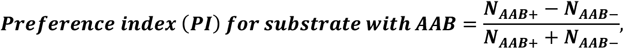

where *N*_AAB+_ and *N*_AAB-_ are the total numbers of eggs on the substrates with acetic acid bacteria (AAB) and the control substrate, respectively.

### 2.4 Feeding assay for *Acetobacter* sp

A binary food choice assay was adapted to analyze feeding site preference by using two different dyes. The chamber for the oviposition assay was used for the feeding assay except that dyed substrates were placed inside the chamber. The substrate was made from 50% diluted apple juice and 1% agar with either blue (brilliant blue FCF, 0.125 mg/mL) or red (sulforhodamine B, 0.1 mg/mL) dyes. The microbial solution and the water control were also dyed with blue or red using the same concentrations as above. The dye colors were randomly switched for each assay.

Flies were starved before the assay in a 50 mL centrifuge tube containing two sheets of Kim-wipe soaked with 3 mL distilled water. The length of starvation time was set differently for each tested group: 24 h for the females of *D. suzukii, D. subpulchrella*, and *D. melanogaster*, 26 h for the females of *D. biarmipes*, 22 h for the males of *D. suzukii*, *D. subpulchrella* and *D. biarmipes*, 20 h for the males of *D. melanogaster*. The temperature was kept at 20 ± 1°C for *D. suzukii* and *D. subpulchrella*, and 25 ± 1°C for *D. melanogaster* and *D. biarmipes*.

After starvation, flies were placed into the feeding chamber without anesthesia and left for 120 min (or 90 min for *D. melanogaster*). Then, the flies were anesthetized by CO2 and were kept at −20°C until the abdomen color was scored under the stereomicroscope.

The feeding preference index (PI) for the substrate inoculated with the microbial solution was calculated with the following formula:

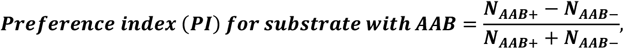

where *N*_AAB+_ and *N*_AAB-_ are the total numbers of flies scored as choosing acetic acid bacteria and control substrate, respectively.

## 3. Results

### 3.1 Oviposition site preferences against *Acetobacter* sp. and *Gluconobacter* sp

In a previous study by our group, we show that in contrast to the females of *D. melanogaster* and *D. biarmipes*, the females of *D. suzukii* did not prefer to lay eggs on substrates inoculated with multiple microbial species collected from other adult flies (Sato et al., 2021). In our current assays, we tested the oviposition preference for a single species of *Acetobacter*, a common constituent of the *Drosophila* gut microbiome (Broderick and Lemaitre 2012; Chandler et al., 2014; Vacchini et al., 2017). A rotten-fruit consumer, *D. melanogaster*, strongly preferred the media with bacterial growth (Figure 1, Supplementary Table S1). Similarly, two tested strains of *D. biarmipes* showed strong preferences for *Acetobacter*. Our current results were consistent with the previous study.

**Figure 1.**
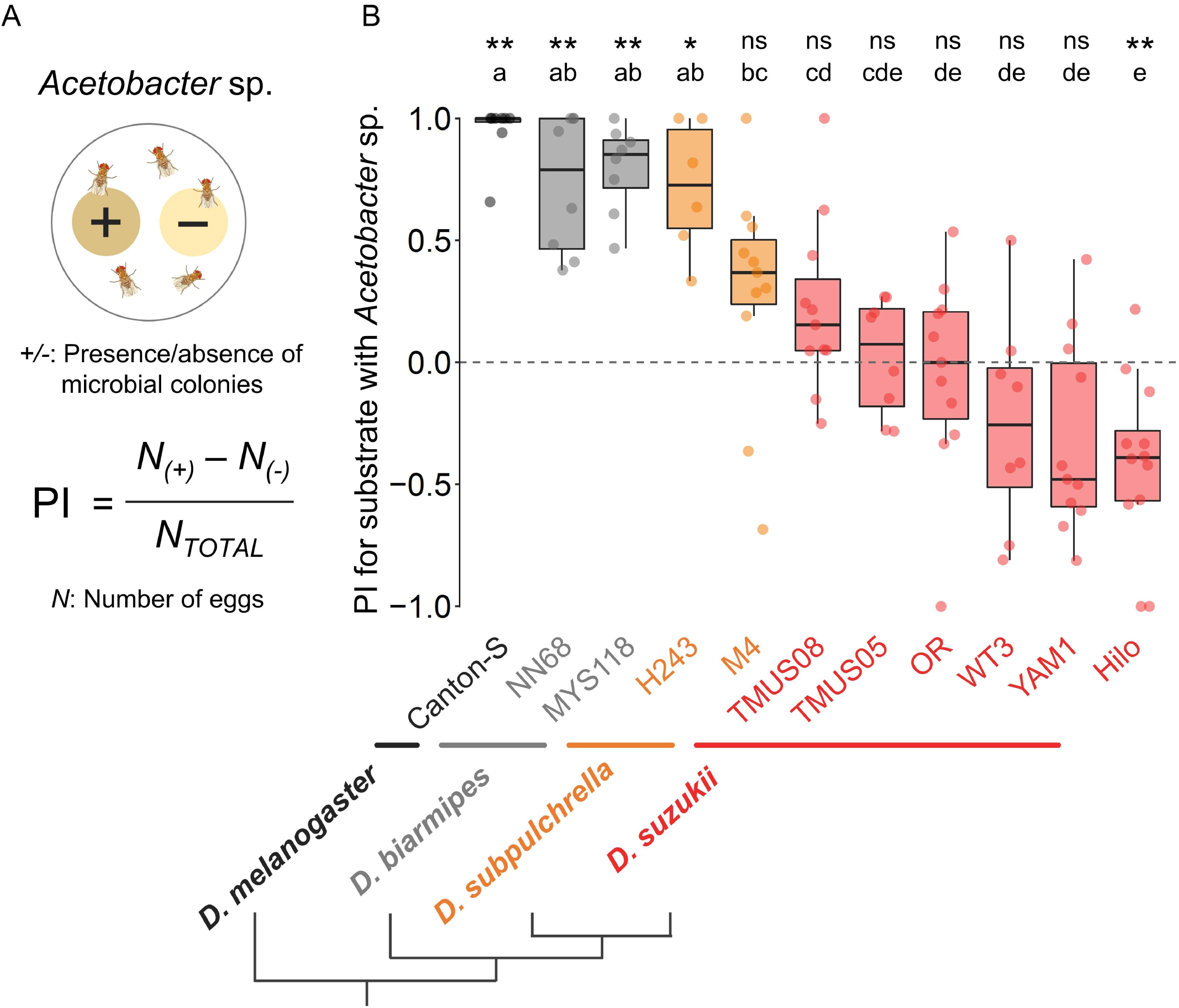
Oviposition site preference quantified as preference index (PI) for *Acetobacter* sp. in *D. suzukii* and its related species. **(A)** Obtaining the PI for oviposition. **(B)** PIs measured using strains from four different species. Results from assays with fewer than 15 eggs on either substrate were excluded from the analyses. Box signifies the upper and lower quartiles and horizontal bar indicates median. Upper and lower whiskers represent maximum and minimum 1.5× interquartile range, respectively. The results from two types of statistical analysis are shown above the graph; the first row indicates the results from two-sided binominal tests assuming an underlying 1:1 proportion (*:*p* < 0.05, **:*p* < 0.01, ns: p ≥ 0.05), and the second row indicates the results from the Kruskal-Wallis tests followed by Dunn’s tests with Benjamin-Hochberg FDR correction (*p* < 0.05).

In contrast, all the strains of *D. suzukii* showed significantly lower PI values compared to the strains of *D. melanogaster* and *D. biarmipes*, suggesting that the preference for *Acetobacter* in *D. suzukii* is distinct from that in *D. melanogaster* and *D. biarmipes*. However, while the Hilo strain avoided *Acetobacter* when choosing the oviposition site, 5 other strains (TMUS05, TMUS08, OR, WT3 and YAM1) did not show any preference or avoidance (Figure 1, Supplementary Table S1). This result implied that there is an intraspecific variation in oviposition site preference for *Acetobacter* in *D. suzukii*.

A sibling species of *D. suzukii, D. subpulchrella*, has diverged after the split from the *D. biarmipes* lineage. The females of this species also have enlarged and serrated ovipositors (Atallah et al., 2014; Muto et al., 2018), however, their tendency to lay eggs into firm substrates or fruits is weaker than that of *D. suzukii* (Atallah et al., 2014; Durkin et al., 2021). Interestingly, while *D. subpulchrella* H243 strain showed a similar *Acetobacter* preference to *D. melanogaster* and *D. biarmipes, D. subpulchrella* M4 strain showed no preference (Figure 1, Supplementary Table S1). There was no significant difference in the PI between *D. subpulchrella* H243 strain and the strains of *D. melanogaster* and *D. biarmipes*. However, the PI of *D. subpulchrella* M4 strain was significantly different from the strains of *D. melanogaster* and *D. biarmipes*, and not different from two of the *D. suzukii* strains (TMUS05 and TMUS08). Therefore, this species has an intermediate degree of preference between *D. suzukii* and *D. melanogaster/D. biarmipes*, and harbors variation within species.

Next, we tested the oviposition site preference for *Gluconobacter*, an acetic acid bacteria family member that is also commonly found in *Drosophila* gut. The assay was conducted using a strain of *D. melanogaster* (Canton-S) and two strains of *D. suzukii* (TMUS08, Hilo), *D*.

4 *subpulchrella* (H243, M4), and *D. biamipes* (NN68, MYS118). Although there was a significant difference between *D. melanogaster* Canton-S and the *D. suzukii* Hilo strain, no significant differences were detected between other strains (Supplementary Figure S1, Supplementary Table S2). These results indicate that the oviposition site preferences for acetic acid bacteria are different between the tested *Acetobacter* and *Gluconobacter* species, exhibiting clearer interspecific divergence for *Acetobacter* than for *Gluconobacter*.

### 3.2 Feeding site preferences against acetic acid bacteria

To our knowledge, binary food choice assays have not been conducted in *Drosophila* species other than *D. melanogaster*. First, to identify the most suitable lengths of time for starvation and feeding assays in females and males of *D. suzukii*, 120-min feeding assays were performed after 24 h of starvation. For *D. suzukii* males, a 22-h starvation period was used because the 24-h period resulted in a high proportion of non-feeding individuals (possibly due to reduced activity caused by excessive starvation) and a high mortality rate (Supplementary Table S3). No preliminary test was performed in *D. subpulchrella*, but the feeding assay could proceed without any problem using the same conditions as for *D. suzukii*.

For *D. biarmipes* and *D. melanogaster*, 90-min feeding assays were initially performed after 24-h starvation as a preliminary test. For *D. melanogaster* males, a 20-h starvation period was used due to the high mortality rate from 24-h tests (Supplementary Table S3). For *D. biarmipes* females, a 26-h starvation period was used because in the 24-h test, the number of deaths during starvation was low while the number of non-feeding individuals during the feeding assay was high, indicating that the flies were inadequately starved (Supplementary Table S4). Because both males and females of *D. biarmipes* did not feed frequently, we performed 120-min feeding assays as with *D. suzukii* and *D. subpulchrella*. Scoring by blue or red abdominal coloration was sufficiently clear in all four assayed species and sexes.

For females of all the tested strains, the median values of the feeding site PIs for *Acetobacter* were positive ranging from 0.13 in *D. suzukii* TMUS08 to 0.64 in *D. melanogaster* Canton-S (Figure 2B). No fixed differences between species were detected and in contrast to the oviposition assay, there was no sign of interspecific divergence among these species. For males, all the tested strains showed no-preference except *D. biarmipes* MYS118 and no significant difference in PI was detected between the strains (Figure 2C).

**Figure 2.**
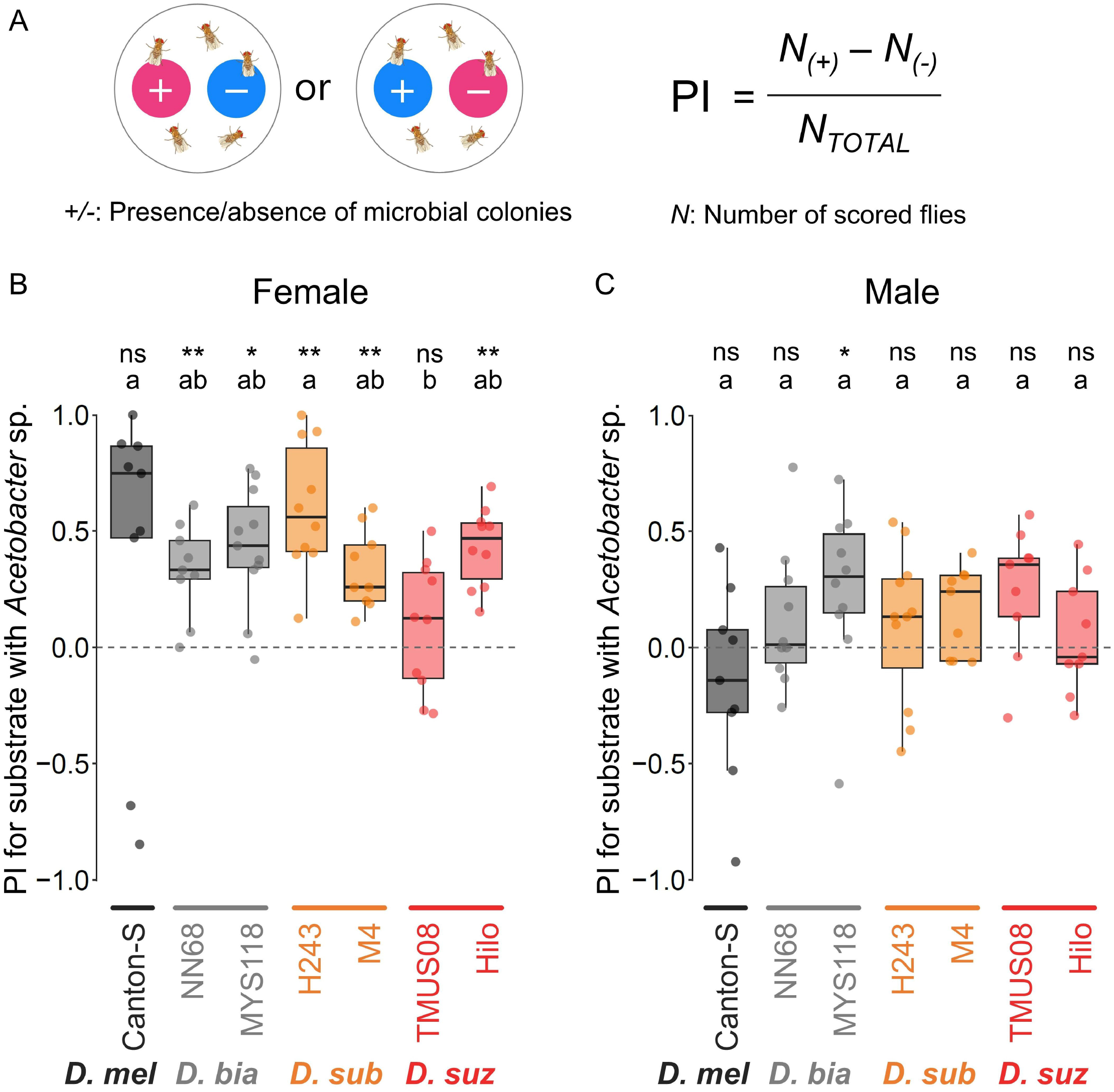
Feeding site preference quantified as preference index (PI) for *Acetobacter* sp. in *D. suzukii* and its related species. **(A)** Obtaining the PI for feeding. **(B)** PIs in females. **(C)** PIs in males. Results from assays with fewer than 80% or 20 scored flies were excluded from the analyses. Box signifies the upper and lower quartiles and horizontal bar indicates median. Upper and lower whiskers represent maximum and minimum 1.5× interquartile range, respectively. The results from two types of statistical analysis are shown above the graph; the first row indicates the results from the two-sided binominal tests assuming an underlying 1:1 proportion (*: *p* < 0.05, ns: *p* ≥ 0.05), and the second row indicates the results of Dunn’s multiple comparisons tests with Benjamin-Hochberg FDR correction (*p* < 0.05).

## 3 Discussion

Ripening fruits provide an open niche for capable fruit fly species to colonize before the resource becomes exploited. The quality of the resource is assessed by different means by the females of *D. suzukii*. For example, firmness, acetic acid concentration, surface curvature, intactness, and the presence of bacteria are among the factors known to affect their oviposition site selection (Atallah et al., 2014; Karageorgi et al., 2017; Kienzle et al., 2020; Durkin et al., 2021; Sato et al., 2021; Akutsu and Matsuo 2022). Among those factors, our oviposition assays focused on the preference for the presence of *Acetobacter* sp. Using multiple strains from each species revealed some intrinsic properties of the shared resource usage among fruit fly species.

Although *D. suzukii* larvae are reported to be more tolerant to low P:C food than other related *Drosophila* species, the intact ripening fruits are not an optimum dietary resource (Silva-Soares et al., 2017). Therefore, the intraspecific variation in the preference for *Acetobacter* growth in our oviposition assay using *D. suzukii* and *D. subpulchrella* could reflect a trade-off between the competitional and nutritional benefits for their offspring when colonizing non-fermenting food. Moreover, the trade-off could be a factor preventing *D. suzukii* from a complete shift to specializing only on ripening fruits.

The interspecific difference in oviposition site preference for *Acetobacter* between *D. suzukii* and *D. biarmipes* was distinct; however, two strains of *D. subpulchrella* represented an intermediate position between the two species. The distribution of *D. suzukii* and *D. subpulchrella* is overlapping and they can be found sympatrically in many localities in Japan (Sasaki and Abe 1993; Takamori et al., 2006; Mitsui et al., 2010). Together with previous studies showing intermediate oviposition characteristics of *D. subpulchrella* between *D. melanogaster* and *D. suzukii* (Atallah et al., 2014; Durkin et al., 2021), our results suggest that the niche separation regarding the oviposition sites between *D. suzukii* and *D. subpulchrella* is not complete.

The oviposition assays in this study revealed that females of most of the tested strains from four different species show a modest preference for media with *Acetobacter* sp. when feeding. The lack of such preference in males from most of the tested strains indicate a higher demand for protein-rich (microbe-rich) food in females than in males (Ribeiro and Dickson 2010; Sun et al., 2017). Also, the comparison of oviposition and feeding site preferences in this study suggest that the interspecific differences in oviposition site preference have evolved independently from the relatively conserved feeding preferences among the tested species.

## 5 Author Contributions

AS, JY and AT conceived the study. AS conducted experiments. AS and AT wrote the manuscript. JY advised on bacteria isolation and culture. All authors edited the manuscript and approved the submitted version.

## 7 Acknowledgments

We thank members of the Takahashi and Yew research groups for helpful discussions and Koichiro Tamura, Seung-Joon Ahn and the Bloomington Drosophila Stock Center for fly stocks.

## 8 Supplementary Material

The supplementary material for this article can be found online.

## 9 Data Availability Statement

The datasets generated for this study can be found in the supplementary materials.

