## Supplementary Figure S1 for "Effect of acetic acid bacteria colonization on oviposition and feeding site choice in *Drosophila suzukii* and its related species"

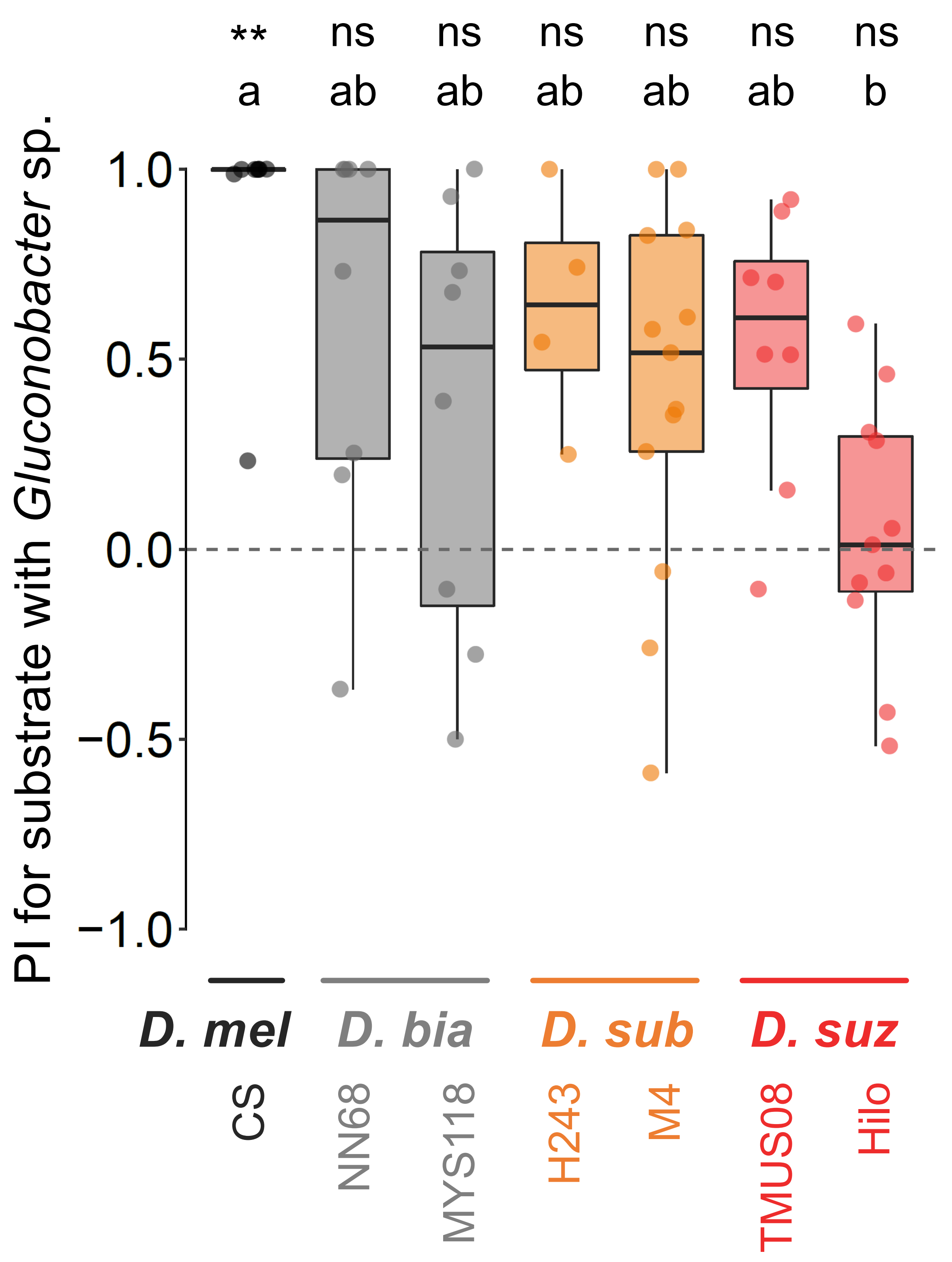


**Figure S1** Oviposition site preferences quantified as preference index (PI) for *Gluconobacter* sp. in *D. suzukii* and its related species. Results from assays with fewer than 15 eggs on either substrate were excluded from the analysis. Box signifies the upper and lower quartiles and horizontal bar indicates median. Upper and lower whiskers represent maximum and minimum 1.5 × interquartile range, respectively. The results from two types of statistical analysis are shown above the graph; the first row indicates the results from two-sided binominal tests assuming an underlying 1:1 proportion (*: $p < 0.05$, **: $p < 0.01$, ns: $p \geq 0.05$), and the second row indicates the results from the Kruskal-Wallis tests followed by Dunn's tests with Benjamin-Hochberg FDR correction ($p < 0.05$).
